# Multivariate consistency of resting-state fMRI connectivity maps acquired on a single individual over 2.5 years, 13 sites and 3 vendors

**DOI:** 10.1101/497743

**Authors:** AmanPreet Badhwar, Yannik Collin-Verreault, Pierre Orban, Sebastian Urchs, Isabelle Chouinard, Jacob Vogel, Olivier Potvin, Simon Duchesne, Pierre Bellec

**Author notes:** **Corresponding Author** Dr. AmanPreet Badhwar, Centre de Recherche, Institut Universitaire de Gériatrie de Montréal, Université de Montréal, Montréal, QC, Canada H3W 1W5.

## Abstract

Studies using resting-state functional magnetic resonance imaging (rsfMRI) are increasingly collecting data at multiple sites in order to speed up recruitment or increase sample size. The main objective of this study was to assess the long-term consistency of rsfMRI connectivity maps derived at multiple sites and vendors using the Canadian Dementia Imaging Protocol (CDIP, www.cdip-pcid.ca). Nine to ten minutes of functional BOLD images were acquired from an adult cognitively healthy volunteer scanned repeatedly at 13 Canadian sites on three scanner makes (General Electric, Philips and Siemens) over the course of 2.5 years. The consistency (spatial Pearson’s correlation) of rsfMRI connectivity maps for seven canonical networks ranged from about 0.4-0.8 (intra-site) to 0.3-0.8 (inter-vendor), with a negligible effect of time. We noted systematic differences in data quality across vendors, which may also explain some of these results. We also pooled the long-term longitudinal data with a single-site, short-term (1 month) data sample acquired on 26 subjects (10 scans per subject), called HNU1. Using randomly selected pairs of scans from each subject, we quantified the ability of a data-driven unsupervised cluster analysis to match two scans of the same subjects. In this “fingerprinting” experiment, we found that scans from the Canadian subject (Csub) could be matched with high accuracy intra-site (>95% for some networks), but that the accuracy decreased substantially for scans drawn from different sites and vendors, while still remaining in the range of accuracies observed in HNU1. Overall, our results demonstrate good multivariate stability of rsfMRI measures over several years, but substantial impact of scanning site and vendors. How detrimental these effects are will depend on the application, yet improving methods for harmonizing multisite analysis is an important area for future work.

**HIGHLIGHTS:**

- Consistency of rsfMRI connectivity over 2.5 years, 13 sites and 3 scanner vendors
- Time elapsed between scans had negligible effect on consistency
- Consistency decreased due to site and vendor differences
- Accuracy of connectivity fingerprints decreased due to site and vendor differences

Graphical Abstract

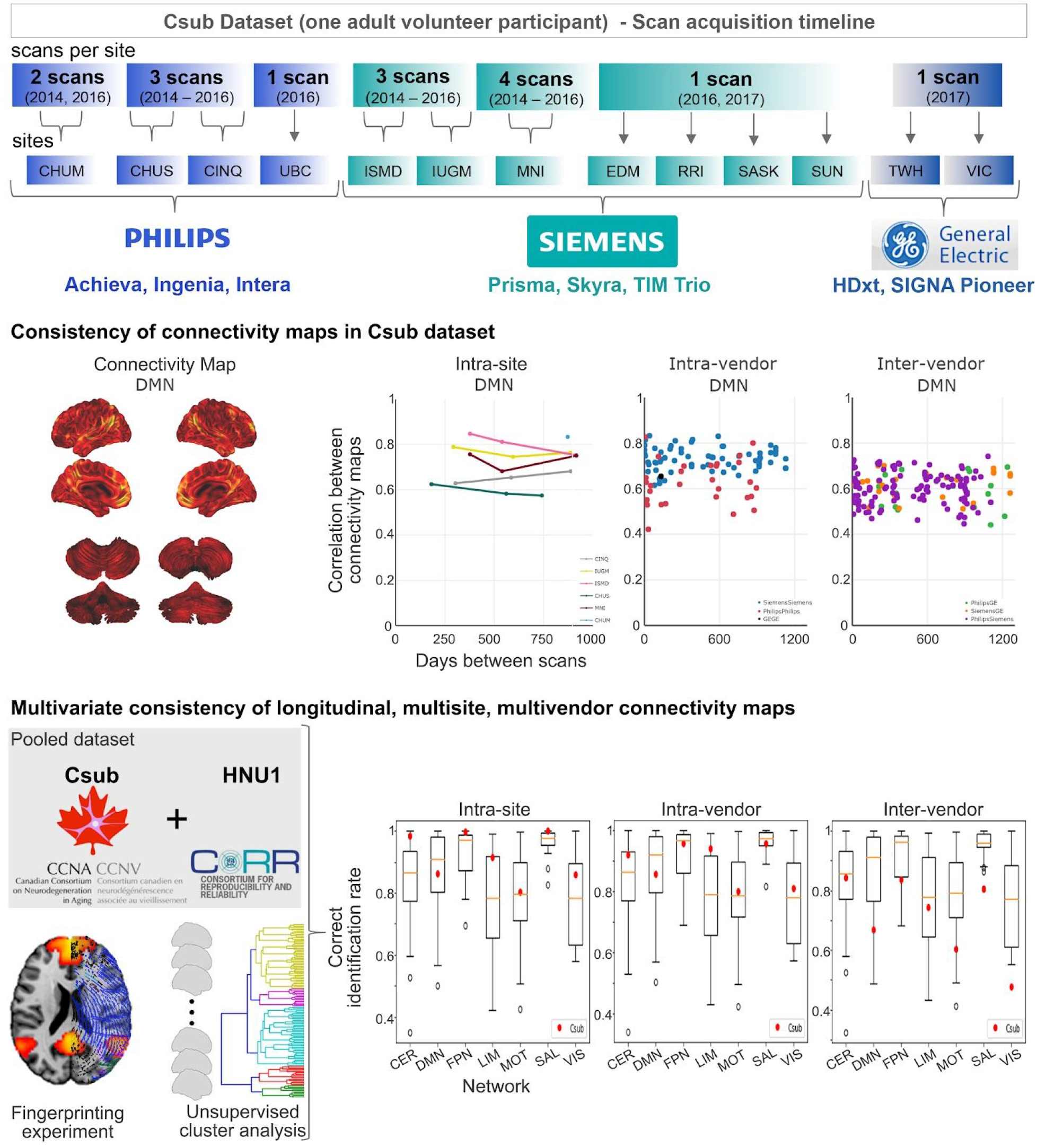

## 1. INTRODUCTION

Paradigm-free (“resting-state”) functional MRI (rsfMRI) can be used to detect spatially distributed functional connectivity networks in health and their alterations in disease (Badhwar et al., 2017; Matthews and Hampshire, 2016). Neuroimaging phenotypes, however, typically exhibit considerable heterogeneity between patients (Dong et al., 2017; Drysdale et al., 2017) and large datasets are needed to achieve sufficient statistical power for reliable detection (Button et al., 2013). Such large patient cohorts frequently surpass the recruitment capacity of single clinical centres. A number of initiatives have pooled multisite data on normal or patient cohorts, such as attention deficit hyperactivity disorder (Brown et al., 2012), autism spectrum disorder (Di Martino et al., 2017; Nielsen et al., 2013), diabetes (Saggar et al., 2017), depression (Drysdale et al., 2017), schizophrenia (Cheng et al., 2015; Skåtun et al., 2017), Alzheimer’s disease (Alzheimer’s Disease Neuroimaging Initiative^1^), population imaging genetics (UK Biobank^2^) and normal brain development (Adolescent Brain Cognitive Development Study or ABCD^3^). However, it is still unclear to what degree the use of multiple scanners introduces additional variance in neuroimaging measures, especially for studies that will collect data for several years. Here we report on the multisite fMRI protocol of a large multisite initiative, lead by the Canadian Consortium for Neurodegeneration in Aging (CCNA^4^), which is recruiting 1600 individuals on the spectrum of age-related dementias over the course of 4 plus years, as well as 660 cognitively normal individuals, at over 30 sites. The CCNA initiative relies on harmonized acquisition parameters set forth in the *Canadian Dementia Imaging Protocol* (CDIP^5^), implementing a series of site qualification, quality control, and assurance procedures.

The main objectives of the present study were to assess the inter-site and longitudinal consistency of rsfMRI measures derived from a single traveling Canadian subject (Csub) scanned repeatedly at several CCNA sites using CDIP. An additional objective was to assess whether the inter-site variance would interfere with a simple machine learning task. We concentrated on fingerprinting (Finn et al., 2015), i.e. identifying paired scans from the same subject in a large multisubject dataset. For this purpose, we pooled the Csub scans with a public dataset featuring multiple retest scans per individual.

The impact of multisite acquisition on rsfMRI connectivity has recently gained attention in a series of studies. Using retrospective rsfMRI data, Yan and colleagues first demonstrated the existence of systematic variations in resting-state connectivity across 18 sites, by contrasting average connectivity patterns of independent groups composed of (mostly) young healthy subjects (Yan et al., 2013). Dansereau et al. further extended this analysis on a subset of eight sites with 3T scanners, showing that average group resting-state network maps could be consistently observed using parcel-based functional connectomes (Dansereau et al., 2017). The authors also reported widespread site effects, present across all resting-state networks, with some sites associated with larger bias than others.

One major limitation of both analyses (Dansereau et al., 2017; Yan et al., 2013) was the reliance on retrospective data, which was a mix of different acquisition parameters (e.g. voxel size, repetition time) as well as scanner make and field strength, all of which may exaggerate the amplitude of site differences. This limitation was addressed by Jovicich and colleagues (Jovicich et al., 2016), who investigated rsfMRI data collected across 13 sites using a harmonized acquisition protocol at 3T on three scanner platforms: Siemens Medical Systems (Siemens), Philips Healthcare (Philips) and General Electric Healthcare (GE). Even with a harmonized protocol, the authors observed significant differences across sites using cross-sectional human volunteer data comprised of independent groups of five participants scanned at each site. This result may partly reflect significant inter-site differences in temporal signal-to-noise ratio (tSNR) maps, observed both on geometric phantoms and volunteer data. This study also scanned each cohort twice over two weeks (median), and demonstrated that retest reliability of connectivity maps was comparable across sites for the major resting-state networks.

A second limitation shared by the multisite rsfMRI studies reviewed thus far was that different participants were recruited at each site, thereby keeping open the possibility that site effects simply reflected differences in participant characteristics. Only a single cohort experiment can unambiguously capture inter-site differences, with the same individual(s) being scanned repeatedly at each site. Three recent studies implemented such an approach. First, Noble and colleagues (Noble et al., 2017a) acquired rsfMRI data on eight participants in two scan sessions separated by 24 hours, and repeated this experiment at eight different sites (all 3T; Siemens and GE scanners) with the same participant cohort and a harmonized protocol. They found that inter-site differences in connectivity measures were substantially explained by the variability of individual within-site measures, especially for short (5 min) acquisition times (Noble et al., 2017a). This conclusion applied to individual region-to-region connectivity, yet multivariate reliability measures from whole-brain connectivity maps were more reliable both within and across sites. In a second, independent study, An and colleagues (An et al., 2017) acquired rsfMRI data on 10 traveling participants in two sessions separated by 30 mins on three scanners, also using a harmonized protocol. Unlike Noble and colleagues, they collected data on Philips scanners in addition to Siemens and GE, but only had one scanner per vendor. Short-term reliability was shown to be better on GE, relative to Siemens and Philips scanners. Unlike Jovicich and colleagues (Jovicich et al., 2016), the authors did not find differences in tSNR ratio across scanner vendors, and there was good reliability of whole brain connectivity maps between scanner vendors. Hawco and colleagues (Hawco et al., 2018) demonstrated that hierarchical clustering can reliably identify functional scans (7 min acquisition times) from four different participants imaged on different scanners across time. Specifically, the authors acquired rsfMRI data on four participants on five different scanners at three sites (all 3T; Siemens and GE scanners), with two subjects scanned long-term (nine scan sessions in 3 years) on one scanner.

A major question left open in the literature is how multisite (>10 sites) acquisitions impact rsfMRI over the long periods of time (years) needed to complete enrolment in large studies such as CCNA. To address this question, we wanted to move beyond traditional measures of consistency for repeated measures (such as intra-class correlation (Fleiss and Cohen, 1973)) because fMRI connectivity maps are high dimensional, multivariate measures, and their primary use case in many instances (e.g. CCNA), will be to serve as features for machine learning prognostic models. Categorical guidelines for interpretation of consistency measures may not translate well in this multivariate predictive context (Cicchetti and Sparrow, 1981; Koo and Li, 2016). Moreover, since functional connectivity maps can act as a ‘fingerprint’ for accurately identifying subjects within a large group (Finn et al., 2015), we selected the accuracy of fingerprinting as a benchmark to assess the consistency of longitudinal fMRI scans. The specific aims and hypotheses of this study were as follows:

a. Evaluate the effect of scanning site, scanner vendor and time elapsed between scans (up to 2.5 years) on the consistency of connectivity maps. Based on the previous literature reviewed above, we hypothesized moderate vendor and site effects. We also hypothesized only a small effect of time: although age effects are detectable in adults, 2 years remain within the error margin of age prediction based on fMRI connectivity (Li et al., 2018).
b. Contrast intra-subject consistency of connectivity maps in a multisite, longitudinal data against intra-subject and inter-subject consistency for a short-term, single site data. Our hypothesis was that site effects would reduce intra-subject consistency (inter-site), but that it would remain higher than inter-subject consistency (intra-site). The rationale for this hypothesis is that the inter-subject differences in brain connectivity are large compared to longitudinal intra-subject differences (Gratton et al., 2018).
c. Evaluate whether the identity of a subject can be reliably identified in the context of multisite, long-term longitudinal data, when pooled with within-site, short-term longitudinal data (fingerprint experiment). In the absence of prior literature, we did not have a specific hypothesis for this aim.

We implemented three experiments to address these aims. We first tested the effect of (a) time, vendor and site, as well as (b) time, vendor, site, tSNR, frame displacement after scrubbing, and volumes remaining after scrubbing using a linear regression analysis of network connectivity maps generated from fMRI data collected on Csub, scanned over 2.5 years at 13 sites using CDIP implemented on one of 3 scanner vendors (results Sections 3.1, 3.2, 3.3 and Supplementary Material, aim a). We then compared the intra-subject consistency in the Csub data with both intra-subject and inter-subject consistency in a public sample^6^ released as part of the Consortium On Resting-state Reproducibility (CORR) (Zuo et al., 2014) and comprised of 30 healthy adults scanned 10 times each over one month (Section 3.4, aim b). Finally, we performed a fingerprinting experiment using scans from the pooled dataset (Section 3.5, aim c), i.e. attempting to match the identity of participants based on pairs of resting-state connectivity maps.

## 2. METHODS

### 2.1 Canadian subject dataset (Csub)

All brain imaging data were acquired from a volunteer Csub: a healthy male with no history of (a) psychiatric and/or neurological illnesses; (b) psychoactive drug usage; or (c) contraindications to MRI. Csub was 42 years old at the start of data collection (2014). In total, the participant underwent 25 scanning sessions at 13 CCNA imaging sites; using scanners from three manufacturers (Philips, Siemens and GE), see Table 1.

**Table 1:**
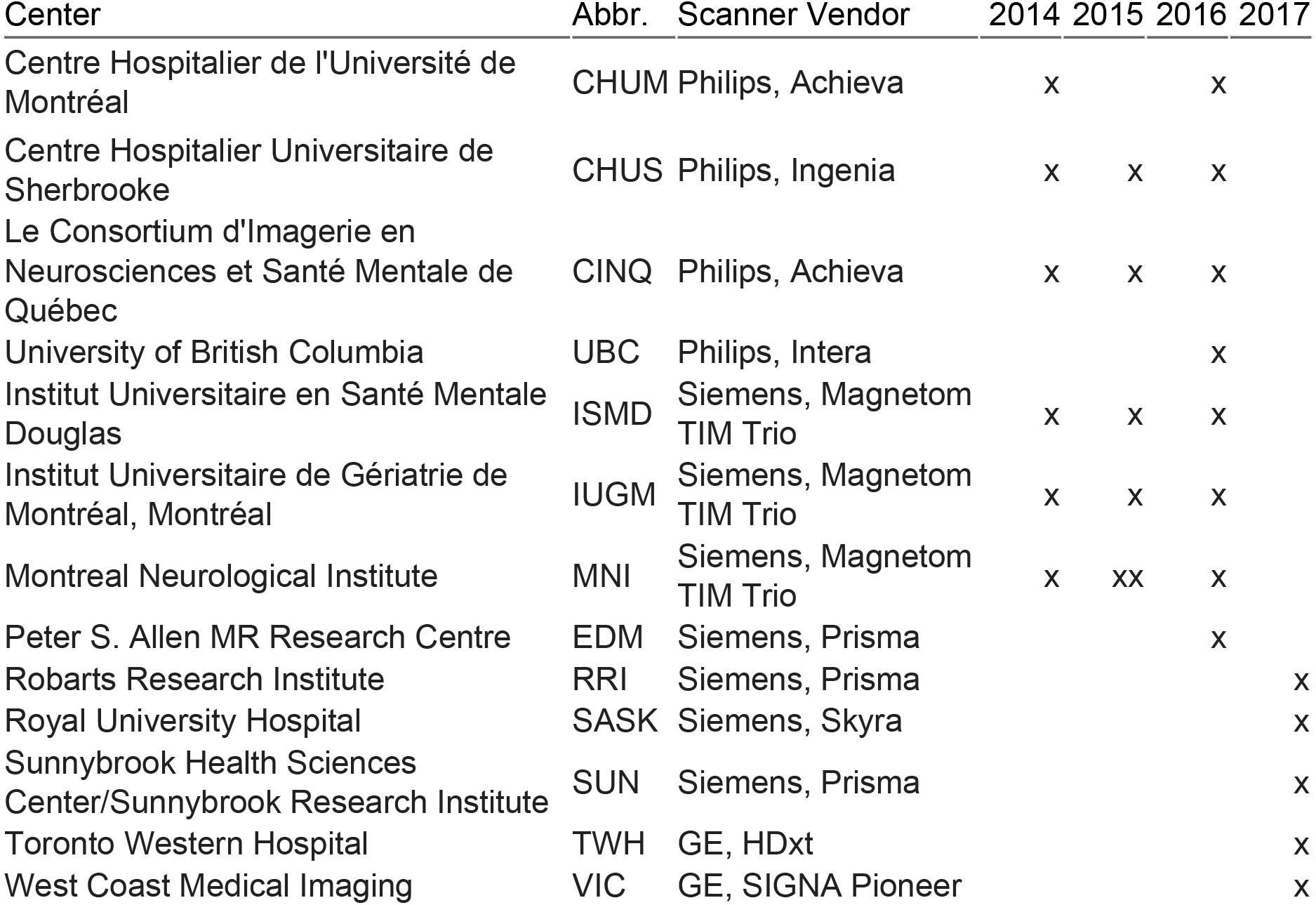
Demographics. The letter “x” in columns 2014-2017 indicate acquisition of rsfMRI and structural scans at the corresponding year.

The data was acquired as part of an ongoing effort to monitor the quality and comparability of MRI data collected across the CCNA imaging network. The schedule of visits did not follow a strict design, with an approximate goal of one visit a year, starting at site qualification. Informed consent was obtained from the subject for the overall study and before every scan session. Due to the multisite nature of the study, ethics approval was obtained from the institutional review board of each participating institution prior to scanning.

Anatomical scans included 3D isotropic T1-weighted (T1w) imaging for assessing fine anatomical detail with high resolution (voxel size = 1.0 × 1.0 × 1.0 mm^3^) and acceleration factor of 2 (Siemens: MP-RAGE; GE: FSPGR; Philips: T1-TFE). Functional T2*-weighted images were obtained using a blood-oxygen-level-dependent (BOLD) sensitive single-shot echo-planar (EPI) sequence. Additional scan parameters are provided in Table 2. During the rsfMRI acquisitions, no specific cognitive tasks were performed, and the participant was instructed to keep his eyes open. No camera or physiological recordings were captured, as these equipments were not available at every site. It should be noted that we excluded the second MNI 2015 intra-session scan (Table 1) from our study, since it was the only intra-session scan acquired. Thereby, data from the remaining 24 scans was used in the study.

**Table 2:**
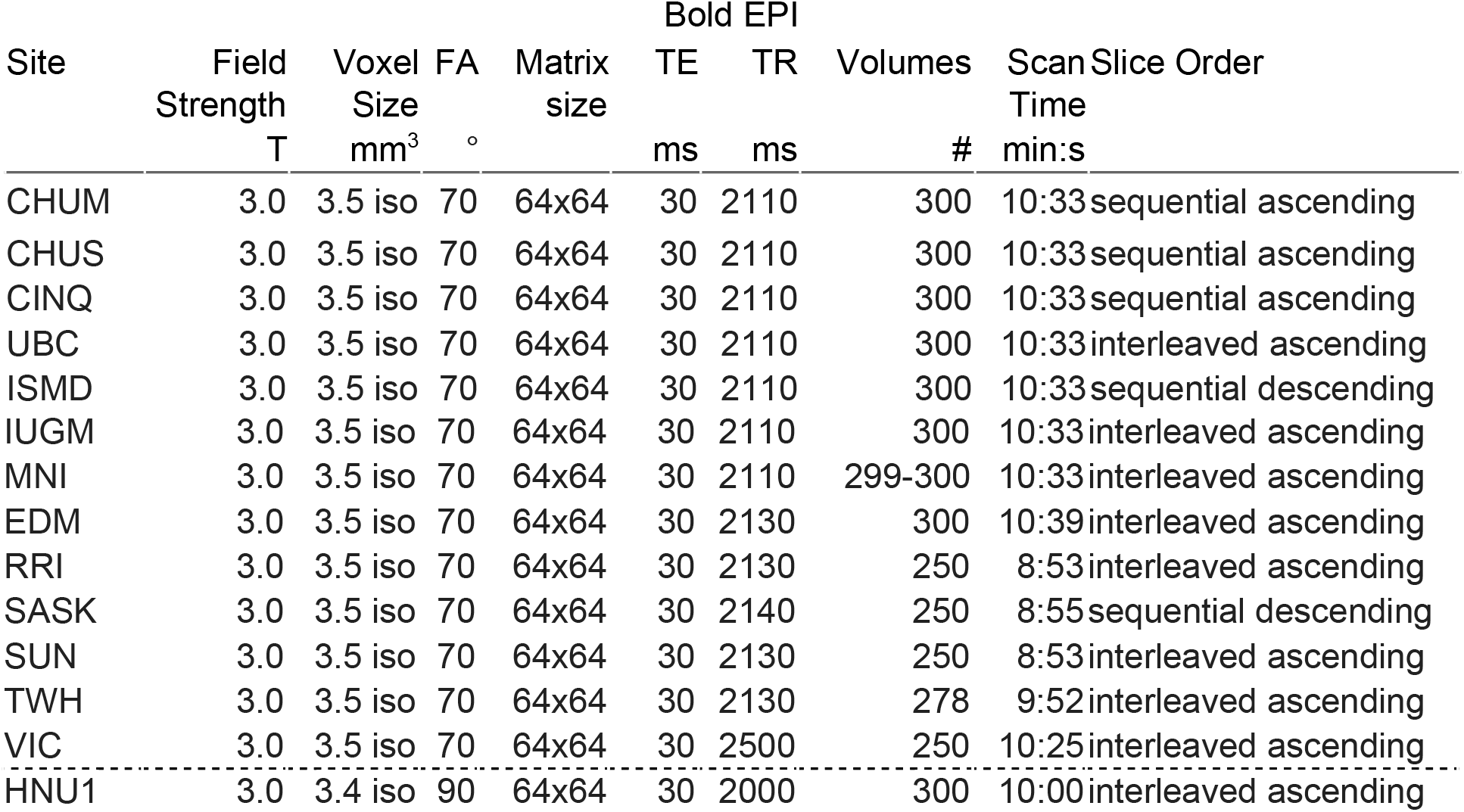
Scan parameters. rsfMRI BOLD EPI scan parameters: Abbreviations: FA, flip angle; ms, millisecond; min:s, minutes and seconds; TE, echo time; TR, repetition time; iso, isometric; T, Tesla; Sites: Full definitions of all the sites have been provided in Table 1.

### 2.2 Hangzhou Normal University dataset (HNU1)

The HNU1 dataset^7^ includes 30 healthy adults 20-30 years of age (mean age 24.4 years), each receiving 10 scans across one month (one scan every three days) on a single 3T GE Discovery MR750 scanner (Zuo et al., 2014). Anatomical scans included 3D isotropic T1w imaging (voxel size = 1.0 × 1.0 × 1.0 mm^3^ and acceleration factor of 2). Functional T2*-weighted images were obtained using a 10 min BOLD-sensitive single-shot EPI sequence^8^. Additional scan parameters are provided in Table 2. During rsfMRI scanning, subjects were presented with a fixation cross and were instructed to keep their eyes open, relax and move as little as possible while observing the fixation cross. Subjects were also instructed not to engage in breath counting or meditation.

### 2.3 Computational environment

The datasets were preprocessed and analyzed using the NeuroImaging Analysis Kit, version 1.1.3 (NIAK-COG^9^, (Bellec et al., 2011)), executed within an Ubuntu 16.0.4 Singularity^10^ container, running GNU Octave11 version 4.2.1, and the MINC toolkit12 version 1.9.15. We also used four Jupyter notebooks that can be executed online via the binder platform^13^, and run in a docker container^14^ built from a public configuration file. Python packages used in the Jupyter notebooks include Numpy (Oliphant, 2006), Pandas (McKinney and Others, 2010), Matplotlib (Hunter, 2007), Scikit-learn (Pedregosa et al., 2011), SciPy (Jones et al., 2016), Seaborn^15^ and StatsModel (Seabold and Perktold, 2010). Interactive plots were generated using Plotly^16^.

### 2.4 Pre-processing of MRI data

With the exception of the T1w scan from UBC, which failed quality control, we averaged all T1w scans (*n*=23) from Csub, following iterative alignment using rigid-body registration. Initially, a brain mask was extracted from a single arbitrary T1w scan (CHUM 2014) using the CIVET pipeline (Ad-Dab’bagh et al., 2006). Remaining T1w scans were then aligned to this reference scan, followed by an averaging of the aligned T1w scans. The averaged image served as the reference scan for the second iteration of alignment and averaging. We performed three such iterations to obtain our final T1w average.

Data from each fMRI scan was corrected for slice timing by linear temporal interpolation. The first three volumes of each fMRI run were discarded to allow the magnetization to reach steady-state. Rigid-body motion was estimated for each time frame, intra-run and inter-run, using CHUM 2014 as an arbitrary reference. Each session was comprised of one fMRI run (see Table 1).

The rigid-body, fMRI-to-T1w and T1w-to-stereotaxic transformations were all combined and used to transform the fMRI images into MNI space at a 3 mm isotropic sampling. The following nuisance covariates were regressed out from the fMRI time series: slow time drifts (basis of discrete cosines with a 0.01 Hz high-pass cut-off), average signals in conservative masks of the white matter and the lateral ventricles, as well as the first principal components (accounting for 95% variance) of the six rigid-body motion parameters and their squares (Giove et al., 2009). The fMRI volumes were finally spatially smoothed with a 6 mm isotropic Gaussian blurring kernel. A more detailed description of the preprocessing pipeline can be found on the NIAK website^17^.

The HNU1 dataset was preprocessed using the NIAK pipeline (Bellec et al., 2011), using the first available structural scan as reference for alignment in stereotaxic space. Four individuals demonstrated subpar alignment of the brain around the meninges. These individuals were excluded from the dataset, and all additional analyses were carried out on the remaining 26 individuals.

### 2.4 Quality control

To minimize artifacts due to excessive motion (Van Dijk et al., 2012), all time frames showing a displacement > 0.5 mm were removed (Power et al., 2012). No scan was excluded due to excessive motion. The number of censored volumes ranged from 0 to 118 time frames, with number of volumes remaining after censoring/scrubbing (n_vols) for generation of connectivity map, ranging from 179 to 297 (std: ±35 volumes, or ±72 seconds taking TR into account). We considered n_vols to be a measure of total available low-motion scan time, a variable known to impact the consistency of connectivity maps (Gordon et al., 2017; Laumann et al., 2015). Frame displacement (FD, scrubbed) values ranged from 0.08 to 0.25. FD is a measure that indexes the movement of the head from one volume to the next. Temporal signal-to-noise ratio (tSNR, raw) values ranged from 58.4 to 108.8. tSNR is an important metric for sensitivity in a given fMRI acquisition protocol and can modulate image intensity over time. Variability in tSNR values are due to both thermal image noise and physiological fluctuations.

Since head motion, scan time, and tSNR are all well known sources of variability, we assessed their relationship with time and scanner vendors. A series of explanatory variables were assembled for a general linear model (GLM) analysis: (1) time between scans, expressed in years and corrected to a zero mean; (2) dummy variables encoding scanner vendors (three covariates: Philips, Siemens minus Philips, and GE minus Philips). A linear mixture of the explanatory variables were adjusted using FD, n_vols and tSNR as dependent variables, separately. We adjusted the significance level of *p* values for multiple comparisons across networks using a Bonferroni correction (family-wise error 0.05, 3 tests, significance threshold *p* <0.05/3, i.e. 0.02). Interpretation of the effect size of FD, n_vols, or tSNR is as follows: the unit of the association is change in consistency (measured by a spatial correlation between maps <= 1) per standard deviation of the specific variable within our sample.

### 2.5 Connectivity maps

Using NIAK’s connectome pipeline^18^, for each rsfMRI scan (from both Csub and HNU1 datasets), we computed voxel-wise connectivity maps associated with each of the seven network templates extracted from a group-level functional brain atlas. Image distortion and signal loss related to magnetic susceptibility artefacts disproportionately impact the frontal cortex, in particular ventrally, as well as temporal cortices, in particular ventromedially. Because of this factor, and possibly other causes, the major brain networks outlined by Yeo-Krienen (Yeo et al., 2011) and others vary substantially by their reliability (Noble et al., 2017b), and their association with disease (Seeley et al., 2009). The networks we used came from the Multiresolution Intrinsic Segmentation Template (MIST) parcellation, which overlaps substantially at this resolution with the Yeo-Krienen atlas (Urchs et al., 2017). The MIST atlas was generated from 200 healthy subjects and consists of nine functional parcellations capturing successively finer levels of spatial detail, of which we used parcellations from resolution seven, consisting of seven commonly used large-scale networks: cerebellar (CER), default-mode (DMN), frontoparietal (FPN), limbic (LIM), motor (MOT), salience (SAL), and visual (VIS). A network connectivity map was obtained per network by computing the Pearson’s correlations between the average time course within the network template and the time course of every voxel in the brain.

### 2.6 Consistency of individual rsfMRI measures within/between sites

For each of the seven rsfMRI networks, a scan by scan similarity (Pearson’s correlation) matrix was generated to summarize the consistency of connectivity maps across the 24 scans in the Csub dataset. A series of explanatory variables were assembled for a general linear model (GLM) analysis: (1) time between scans, expressed in years and corrected to a zero mean; (2) dummy variables encoding intra-vendor comparisons (three covariates: GE, Siemens and Philips); (3) dummy variables encoding intra-site comparisons (six covariates: CHUM, CHUS, CINQ, ISDM, IUGM, MNI; the other sites did not have multiple retest data available). An intercept was also added to the model, which, in combination with the other covariates of the model, captured the average consistency for comparisons across sites from different vendors. A linear mixture of the explanatory variables were adjusted on the inter-scan consistency measures (dependent variable) using ordinary least squares, for each network separately. For each network, we tested the significance of the effect of inter-vendor (t-test), intra-vendor (*F* test testing the combined effect of the three intra-vendor covariates), intra-site (*F* test testing the combined effect of the six intra-site covariates) and time (*t*-test). We adjusted the significance level of *p* values for multiple comparisons across networks using a Bonferroni correction (family-wise error 0.05, 7 networks × 3 tests = 21 total tests; significance threshold *p* <0.05/21, i.e. 0.002). We also examined the effects of each individual variable to assess which vendors and sites drove the significance of tests.

We also performed a second analysis that included the initial explanatory variables plus three additional variables: FD as a metric of head motion; n_vols as a metric of total available low-motion scan time; and tSNR as a metric of fMRI data quality. We tested the impact of these additional variables using t-tests, and adjusted the significance level of *p* values for multiple comparisons across networks using a Bonferroni correction (family-wise error 0.05, 7 networks × 6 tests = 42 total tests; significance threshold *p* <0.05/42, i.e. 0.001).

### 2.7 Consistency of rsfMRI measures within/between subjects

For each HNU1 subject and each network, we computed the average (and standard deviation) for the intra-subject consistency for all pairs with the 10 available scans. We also computed the average inter-subject consistency across all scans from different subjects, both within HNU1, and between HNU1 and Csub. To further statistically compare these consistency values, we implemented a single GLM analysis, in which the dependent variable was the measures of inter-scan consistency, and the explanatory variables included a series of dummy variables encoding separately the intra-subject comparisons (Csub and 26 HNU1 subjects), one dummy variable encoding comparisons between Csub and HNU1 subjects, and one dummy variable encoding inter-subject comparisons in HNU1. A series of t-test were derived from the following contrasts: intra-subject in Csub vs intra-subject in HNU1, intra-subject in Csub vs inter-subject in HNU1, inter-subject Csub/HNU1 vs inter-subject HNU1, and intra-subject HNU1 vs inter-subject HNU1.

### 2.8 Fingerprinting of HNU1 participants and Csub

We assessed the ability of a simple data-driven cluster analysis to recover the identity of subjects based on connectivity maps of a single network, mixing the Csub single subject with the HNU1 subjects. A fingerprinting experiment consisted of the following steps: (i) for each subject, randomly select two scans out of all available scans (at least 10); (ii) assemble an inter-scan similarity matrix, using only the selected scans for all subjects; (iii) apply a hierarchical clustering on this similarity matrix (Ward’s criterion); (iv) group the scans into as many clusters as there are subjects, based on the hierarchy; (v) for each subject, the fingerprinting experiment is considered successful if the two scans of this subject constitute a cluster.

The fingerprinting procedure was repeated *B*=10000 times using random scan selections, and independently for each network. The average accuracy of fingerprinting for a given subject and network was derived as the proportion of successful fingerprinting repetitions. In addition, we ran three different types of experiments. First, the pair of scans for Csub were drawn from the same site (intra-site fingerprinting experiment), second, the pair of scans for Csub were drawn from different sites, but the same vendor (intra-vendor fingerprinting experiment), and third, the pair of scans for Csub were drawn from different sites and different vendors (inter-vendor fingerprinting experiment).

### 2.9 Data Records

Scripts used in this study are available on Github^19^, as well as archived on zenodo^20^. The four Jupyter notebooks used in this study (graphs, stats_repro, stats_tsnr_time_motion and stats_fingerprinting) can be executed online via the binder platform^21^. We have also made available on Github and zenodo two interactive dashboards containing (1) connectivity maps used to assess long-term consistency in Csub rsfMRI measures, and (2) connectivity maps from Csub and HNU1 datasets. Provided in each dashboard is the individual connectivity map per network, the average connectivity map per network, and the MIST parcellation at scale 7.

## 3 RESULTS

### 3.1. Relationship of head motion, scan-time and tSNR with time and scanner vendors

There was no significant (p<0.02) impact of time on either FD (scrubbed), n_vols or tSNR (Table 3).

**Table 3:** Relationship of head motion, scan time, and tSNR with time and scanner vendors. (*) indicates family-wise error < 0.05 (Bonferroni corrected for multiple comparisons, adjusted threshold p <0.02).

|  | Relationship with FD |  |  | Relationship with n_vols |  |  | Relationship with tSNR |  |  |
| --- | --- | --- | --- | --- | --- | --- | --- | --- | --- |
|  | coef | std err | p value | coef | std err | p value | coef | std err | p value |
| Philips | 1.1398 | 0.186 | *0.000 | -0.6790 | 0.292 | 0.031 | 0.7339 | 0.309 | 0.028 |
| time | 0.2195 | 0.106 | 0.052 | -0.1404 | 0.167 | 0.411 | 0.1359 | 0.177 | 0.451 |
| Siemens<br>minus Philips | -1.8427 | 0.240 | *0.000 | 1.2311 | 0.377 | *0.004 | -1.1898 | 0.398 | *0.007 |
| GE<br>minus Philips | -1.7000 | 0.473 | *0.002 | 0.1456 | 0.744 | 0.847 | -1.0733 | 0.787 | 0.188 |

<sup>19</sup> https://github.com/SIMEXP/cdip_human_phantom
<sup>20</sup> https://doi.org/10.5281/zenodo.3350885
<sup>21</sup> https://mybinder.org/v2/gh/SIMEXP/cdip_human_phantom/master

There was however significant (p<0.02) differences in average FD between all three scanner vendors (Table 3). Philips and Siemens demonstrated a significant difference in average n_vols and tSNR (Table 3). Specifically, Philips scans had higher motion levels than Siemens scans, which resulted in lower number of volumes after scrubbing. Philips scans also had higher tSNR than Siemens scans.

### 3.2 Connectivity maps

The key regions of all 7 networks were clearly identifiable at every session, as illustrated for DMN connectivity (Figure 1). Random fluctuations were also apparent, sometimes with strong shifts in global connectivity values (see for example IUGM vs EDM in Figure 1).

**Figure 1:**
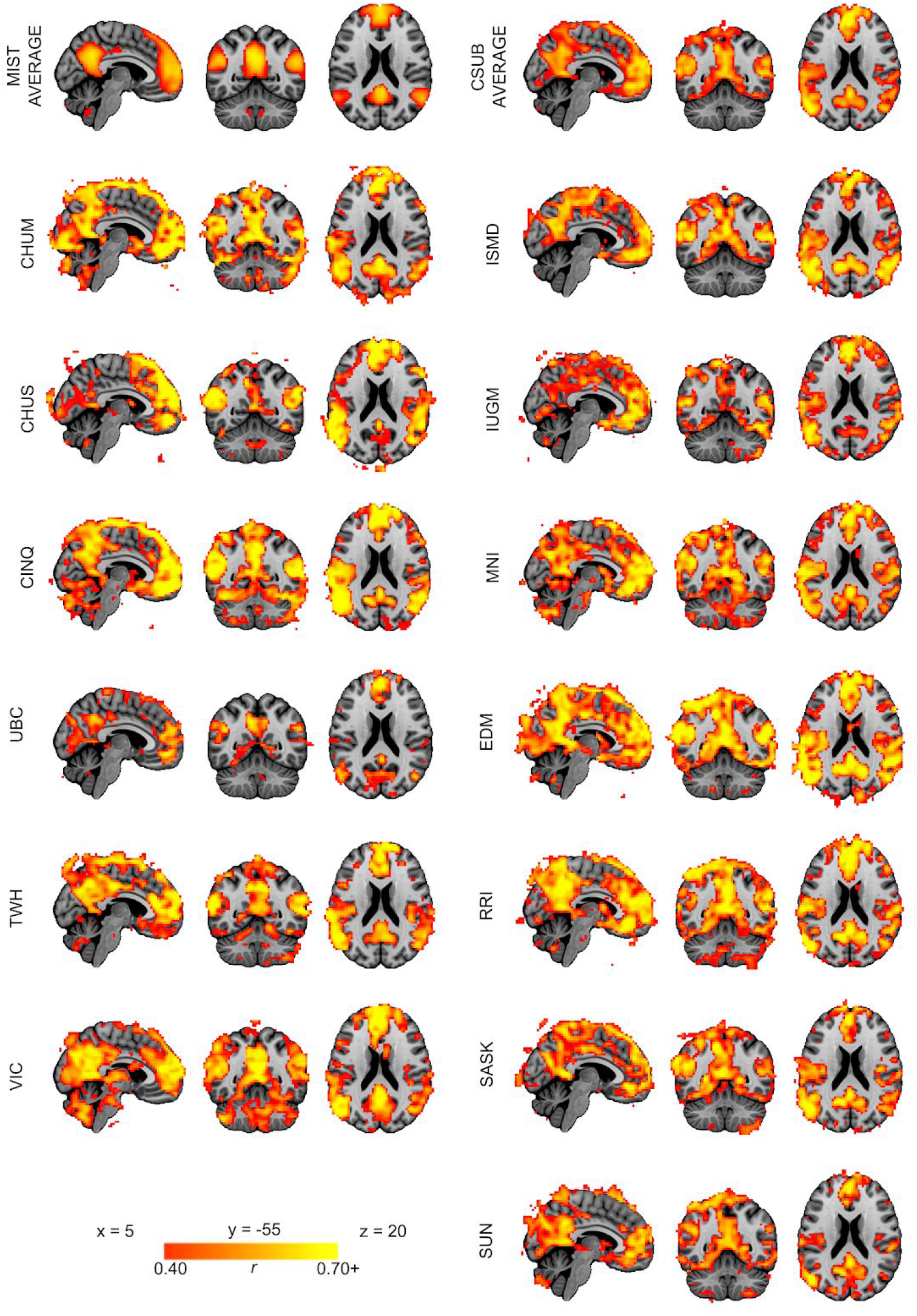
Default-mode network connectivity. The connectivity map for the DMN of each site is included. For sites with multiple sessions, only the most recent is shown. Brighter colours (orange-yellow) in the connectivity maps indicate stronger connectivity strength (higher Pearson r correlation). Maps are superimposed onto the anatomic International Consortium for Brain Mapping (ICBM) 152 template. Top left map is a DMN average of the MIST parcellation dataset. Top right map is a DMN average of all 13 sites from the Csub dataset. Full names of all the sites have been provided in Table 1.

We then quantified the consistency of rsfMRI maps generated at different sessions using the Pearson’s correlation coefficient as a measure of spatial similarity. We selected this measure as it is invariant to the shifts in mean and variance we noted above

### 3.3 Consistency of individual fMRI measures within/between sites

The consistency of maps generated with rsfMRI data acquired on scanners from different vendors ranged from 0.57 ± 0.01 (limbic network) to 0.64 ± 0.01 (visual network), see Table 4. There was no substantial (or significant) effect of time between scanning sessions on consistency between maps. The estimated yearly rate of change in consistency (measured on a spatial correlation scale from −1 to 1) ranged from 4.22e-4, p = 0.26 (limbic network) to −3.98e-3, p = 0.28 (visual network), see Table 4, Figure 2.

**Table 4:**
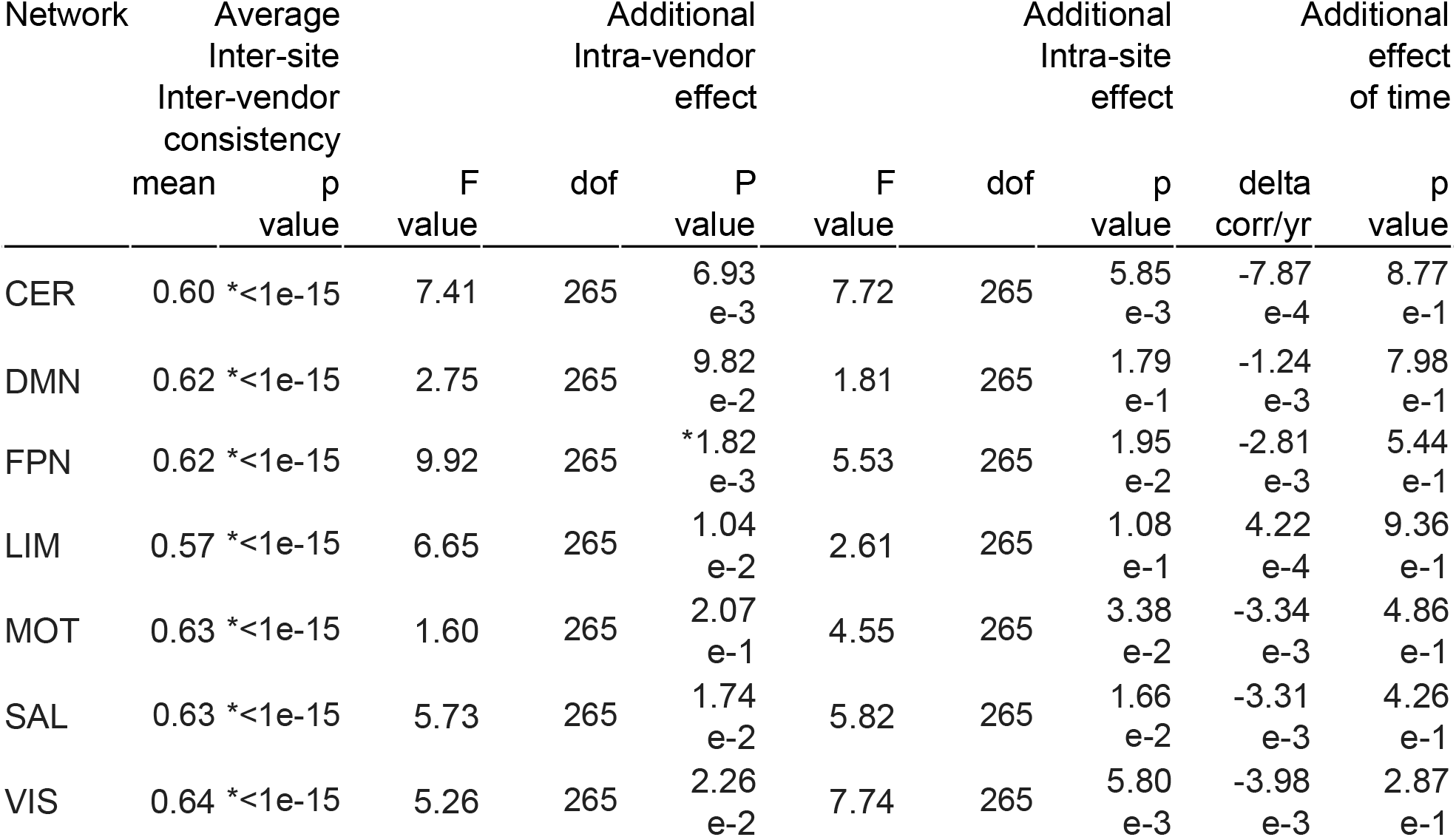
Effect of vendor, site and time on consistency of rsfMRI connectivity measures. (*) indicates family-wise error < 0.05 (Bonferroni corrected for multiple comparisons across networks, adjusted threshold p<0.002). Abbreviations: Networks: CER, cerebellar; DMN, default mode; FPN, frontoparietal; LIM, limbic; MOT, motor; SAL, salience; VIS, visual.

| Network | Average Inter-site Inter-vendor consistency |  | Additional Intra-vendor effect |  |  | Additional Intra-site effect |  |  | Additional effect of time |  |
| --- | --- | --- | --- | --- | --- | --- | --- | --- | --- | --- |
|  | mean | p value | F value | dof | P value | F value | dof | p value | delta corr/yr | p value |
| CER | 0.60 | $<1e-15$ | 7.41 | 265 | $6.93e-3$ | 7.72 | 265 | $5.85e-3$ | $-7.87e-4$ | $8.77e-1$ |
| DMN | 0.62 | $<1e-15$ | 2.75 | 265 | $9.82e-2$ | 1.81 | 265 | $1.79e-1$ | $-1.24e-3$ | $7.98e-1$ |
| FPN | 0.62 | $<1e-15$ | 9.92 | 265 | $*1.82e-3$ | 5.53 | 265 | $1.95e-2$ | $-2.81e-3$ | $5.44e-1$ |
| LIM | 0.57 | $<1e-15$ | 6.65 | 265 | $1.04e-2$ | 2.61 | 265 | $1.08e-1$ | $4.22e-4$ | $9.36e-1$ |
| MOT | 0.63 | $<1e-15$ | 1.60 | 265 | $2.07e-1$ | 4.55 | 265 | $3.38e-2$ | $-3.34e-3$ | $4.86e-1$ |
| SAL | 0.63 | $<1e-15$ | 5.73 | 265 | $1.74e-2$ | 5.82 | 265 | $1.66e-2$ | $-3.31e-3$ | $4.26e-1$ |
| VIS | 0.64 | $<1e-15$ | 5.26 | 265 | $2.26e-2$ | 7.74 | 265 | $5.80e-3$ | $-3.98e-3$ | $2.87e-1$ |

**Figure 2:**
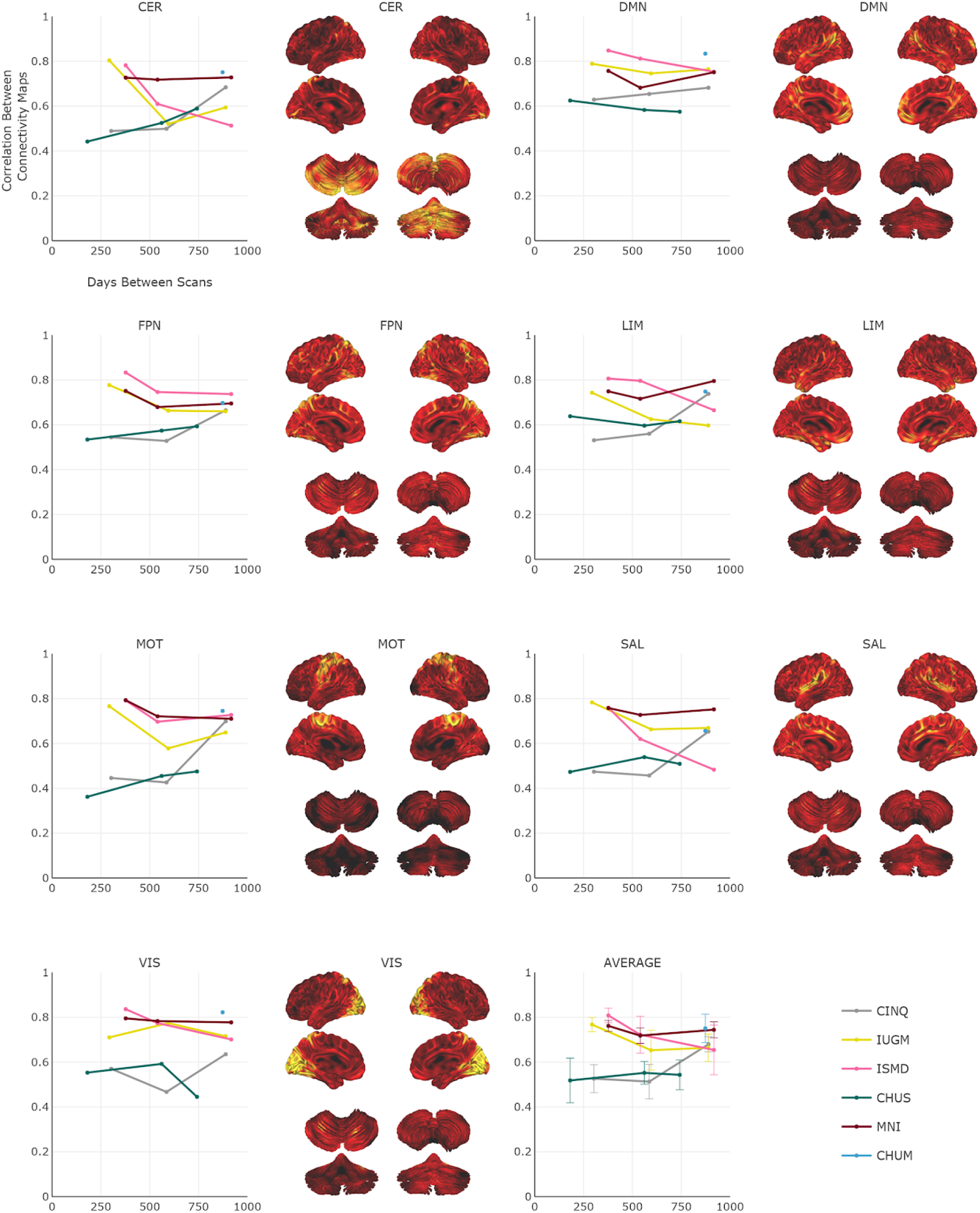
Intra-site consistency over time. Per network, the intra-site consistency (Pearson’s correlation r) over time (range: 0 to 917 days) for the six sites. Longitudinal data are presented as line plots on the left, and the average connectivity map is provided on the right. Brighter colours (orange-yellow) in the connectivity maps indicate stronger connectivity strength. Maps are superimposed onto the anatomic International Consortium for Brain Mapping (ICBM) 152 template and the SPM2_MNI aligned cerebellum surface (Van Essen et al. 2004). The average consistency across all networks are also shown. Abbreviations: Networks: CER, cerebellar; DMN, default mode; FPN, frontoparietal; LIM, limbic; MOT, motor; SAL, salience; VIS, visual. Sites: Full names of all the sites have been provided in Table 1. Note: Interactive graphs are provided in the “graphs” Jupyter notebook.

There was a significant effect of vendors in one network (frontoparietal), with trends (p<0.05 uncorrected) in four others (cerebellar, limbic, salience and visual), see Table 4 and Figure 3. This suggests that, for this network, inter-site, intra-vendor consistency was significantly different from inter-site, inter-vendor consistency. The effect was driven by Siemens scanners, with markedly higher consistency in all seven networks (see Supplementary Material Table S1).

**Figure 3:**
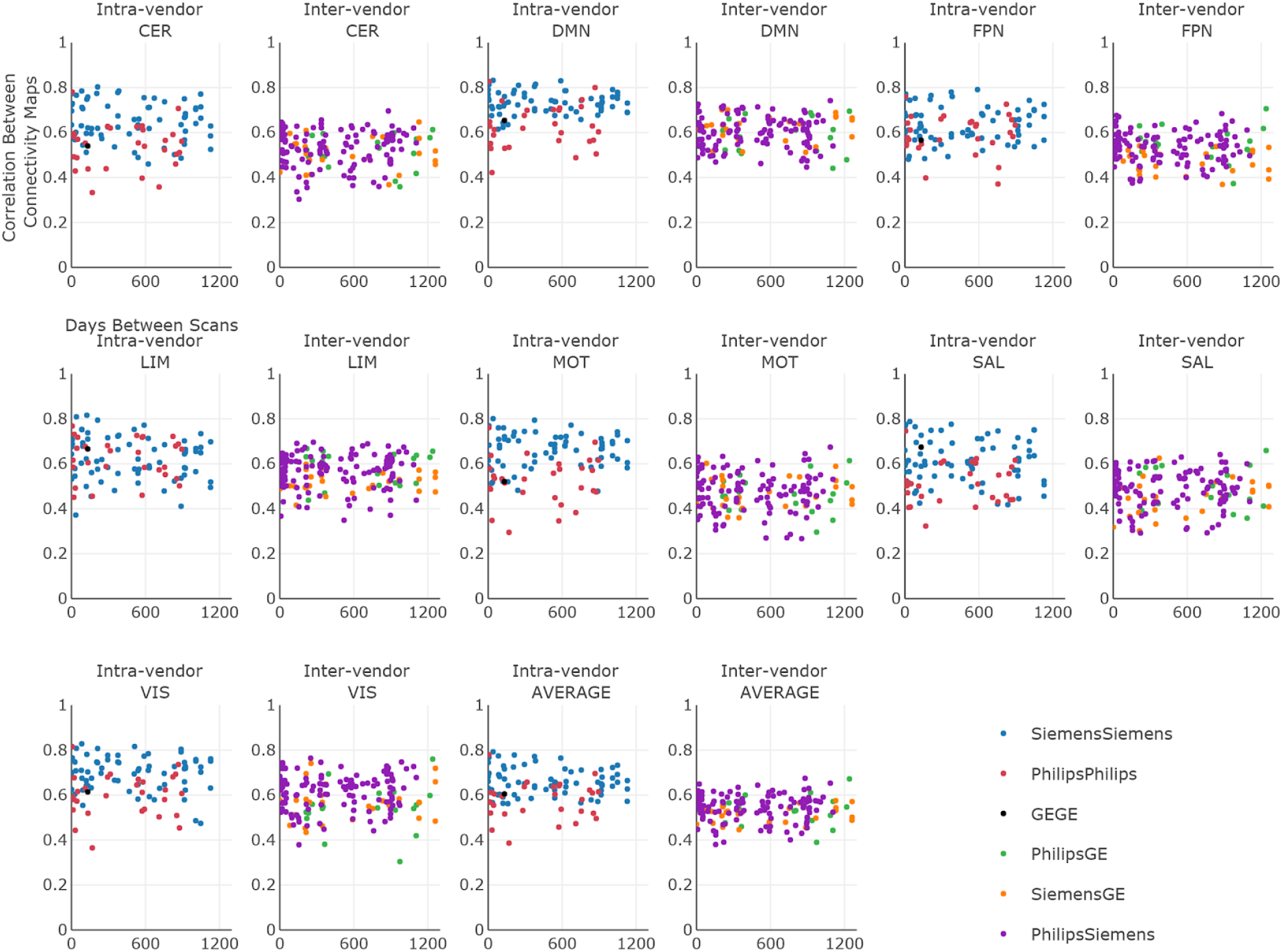
Intra-vendor and inter-vendor consistency over time. Per network, the intra- and inter-vendor consistency over time (ranging from 0 to 1240 days) for GE, Philips and Siemens are presented as scatter plots. Intra- and inter-vendor average consistency across all networks are also shown. Abbreviations: Networks: CER, cerebellar; DMN, default mode; FPN, frontoparietal; LIM, limbic; MOT, motor; SAL, salience; VIS, visual. Note: Interactive graphs are provided in the “graphs” Jupyter notebook.

Intra-site consistency was not significantly higher than inter-site, inter-vendor consistency in any of the seven networks, see Table 4, though trends (p<0.05 uncorrected) were observed in five networks (cerebellar, frontoparietal, motor, salience and visual). The intra-site effects on consistency were highly heterogeneous, with some sites showing very small effects (e.g. cerebellar network: CINQ, difference in consistency 0.0037, p=0.942), while others were markedly different (e.g. cerebellar network: MNI, difference in consistency 0.1536, p=0.002), see Supplementary Material Table S1.

The second consistency analysis that also included tSNR, FD and n_vols as dummy variables was more challenging to interpret since the three variables are related to the inter-vendor variables. Therefore, the additional impact of inter-vendor effects on consistency was now distributed over these three variables (See Supplementary Material Tables S2 and S3).

### 3.4 Consistency of fMRI measures within/between subjects

We evaluated both intra- and inter-subject consistency in HNU1 subjects for all seven networks (Table 5 and Supplementary Material Figure S1).

**Table 5:** Consistency of intra-subject and inter-subject connectivity measures. Abbreviations: Networks: CER, cerebellar; DMN, default mode; FPN, frontoparietal; LIM, limbic; MOT, motor; SAL, salience; VIS, visual.

| Network | HNU1 | HNU1 | Csub | Csub vs HNU1 |
| --- | --- | --- | --- | --- |
|  | intra-subject | inter-subject | intra-subject | inter-subject |
| CER | $0.64 \pm 0.10$ | $0.42 \pm 0.08$ | $0.59 \pm 0.09$ | $0.37 \pm 0.07$ |
| DMN | $0.73 \pm 0.13$ | $0.48 \pm 0.10$ | $0.56 \pm 0.10$ | $0.33 \pm 0.07$ |
| FPN | $0.68 \pm 0.12$ | $0.39 \pm 0.09$ | $0.53 \pm 0.11$ | $0.26 \pm 0.09$ |
| LIM | $0.60 \pm 0.12$ | $0.37 \pm 0.09$ | $0.53 \pm 0.12$ | $0.30 \pm 0.07$ |
| MOT | $0.72 \pm 0.10$ | $0.50 \pm 0.09$ | $0.63 \pm 0.10$ | $0.43 \pm 0.10$ |
| SAL | $0.73 \pm 0.11$ | $0.47 \pm 0.08$ | $0.57 \pm 0.09$ | $0.35 \pm 0.07$ |
| VIS | $0.75 \pm 0.10$ | $0.56 \pm 0.09$ | $0.65 \pm 0.09$ | $0.48 \pm 0.07$ |

Average intra-subject consistency ranged from 0.60 ± 0.12 (limbic network) to 0.75 ± 0.10 (visual network). Intra-subject consistency in HNU1 was higher than the average intra-Csub consistency across all networks (e.g. 0.73 ± 0.13 in HNU1 vs 0.56 ± 0.10 in Csub, for the DMN, all tests *p*<10^-15). Inter-subject consistency in HNU1 was lower than intra-subject consistency, both HNU1 and Csub (e.g. inter-subject of 0.48 ± 0.10 in HNU1, versus intra-subject of 0.73 ± 0.13 in HNU1 and intra-subject of 0.56 ± 0.10 in Csub, for the DMN, all tests *p*<10^-15). The consistency between HNU1 participants and Csub was also lower than the consistency between HNU1 participants across all networks (e.g. inter-subject of 0.48 ± 0.10 in HNU1 vs 0.33 ± 0.0 for Csub vs HNU1, in the DMN, all tests *p*<10^-15). Overall, the site effects present in Csub scans seemed to decrease both the intra- and inter-subject consistency, compared to monosite HNU1 data. Yet, intra-subject Csub consistency remained higher than inter-subject HNU1 consistency.

### 3.5 Fingerprinting of HNU1 participants and Csub

We ran fingerprinting experiments by mixing 2 random scans of Csub with 2 random scans for each of the HNU1 participants, and using an unsupervised cluster analysis to determine whether the two scans of a subject would be clustered together. Experiments were replicated using pairs of scan selected either intra-site, intra-vendor or inter-vendor for Csub. For all three experiments, the highest fingerprinting accuracy was reached for the salience and frontoparietal networks, with about 90+% successful identification on median across HNU1 subjects (Figure 4). Some networks reached lower median accuracy (<80%), but still much higher than chance level (1/26 subjects ~ 4%).

**Figure 4:**
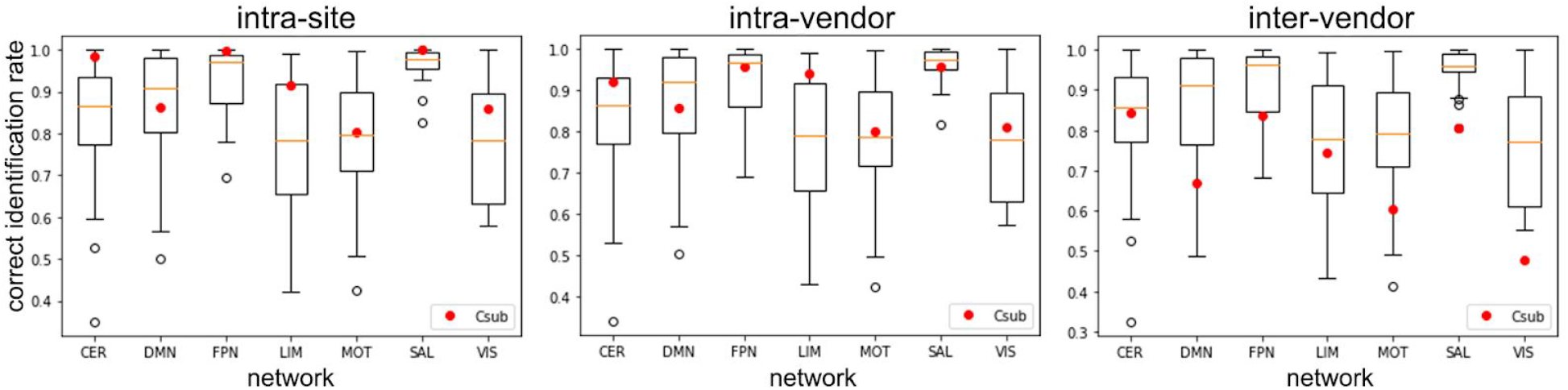
Fingerprinting accuracy. Distribution of the correct identification rate per network for the 26 HNU1 subjects. The red circle indicates the average identification accuracy of Csub per network in the intra-site, intra-vendor, and inter-vendor experiments.

Accuracy observed for Csub was very high for the intra-site experiments, reaching almost perfect accuracy in the cerebellar, frontoparietal and salience networks, and was generally on the higher end of the HNU1 distribution, with the exception of the default mode network. There was a clear decrease in accuracy (over 0.2 or even 0.3 in some networks) when moving from intra-site to intra- and inter-vendor. In particular, Csub fell in the lower half of the HNU1 distribution for the default mode, frontoparietal and salience networks for the intra-vendor experiment; and then in the lower end of the HNU1 distribution for all seven networks for the inter-vendor experiment. Csub actually fell outside of the HNU1 distribution only for the VIS network in the inter-vendor experiment.

## 4. DISCUSSION

### 4.1 Consistency in rsfMRI connectivity measures within/between sites

In the present study, we assessed the consistency of rsfMRI connectivity measures in a single participant, scanned at 13 CDIP-compliant sites. We report consistencies of 0.53 ± 0.11 (frontoparietal network) to 0.65 ± 0.09 (visual network) for connectivity maps generated from data obtained at different sites, scanner vendors and time points separated by a wide range of durations, from 0 to 1262 days apart. We found significant effects of scanner vendors, although only one vendor (Siemens) was associated with substantial effect of higher consistency. The finding that Siemens scanners have more consistent maps than Philips scanners is in agreement with the report of An and colleagues (An et al., 2017). We did not replicate the excellent consistency of GE scanners, but we had only one inter-GE scanner comparison available in the Csub sample. Such inter-vendor differences in consistency may be due to factors such as scanner drift (Friedman and Glover, 2006) and smoothness of the raw images produced (Friedman et al., 2006).

With regards to our intra-site consistency metric, Laumann et al. (Laumann et al., 2015) reported a very similar metric to the one used: correlation of connectomes between test and retest, as a function of time used to estimate the connectome (while we worked on voxel-based maps). Their graph is hard to read precisely for small amounts of time, but 10 minutes seems associated with a correlation of at least 0.8, which is exactly what we observed in our sample intra-site. Moreover, our observation that site effects were often very small is in line with the observation of Noble and colleagues (Noble et al., 2017a). This study found that, especially with short time series (less than 10 min), the physiological variability of resting-state measures dominates the scanner variations. Our report extends upon this analysis with longer longitudinal follow up (years instead of weeks), more sites (13 instead of 8), and more vendors (three instead of two).

### 4.2 Consistency within/between subjects

We found in the HNU1 data that the similarity of maps generated between different individuals is much lower (e.g. 0.48 ± 0.10 for DMN) than the similarity observed intra-subject (e.g. 0.73 ± 0.13 for DMN) in the same sample, as well as intra-subject in our Csub participant (e.g. 0.56 ± 0.10 for DMN). This last consistency value is an average across many scan sites, vendors, and inter-scan intervals and, consequently, the intra-subject Csub consistency was lower than in HNU1 (on average across subjects). The observation suggests that, even with multisite, long-term longitudinal data and relatively short scan duration (about 10 minutes for both Csub and HNU1), it may be possible to implement reliable “brain fingerprinting”. Potential feasibility of fingerprinting was also reinforced by the observation that comparisons between Csub and HNU1 participants were lower on average (e.g. 0.33 ± 0.07 for DMN) than inter-subject comparisons in HNU1 (e.g. 0.48 ± 0.10 for DMN).

### 4.3 Fingerprinting

We found that it was possible to fingerprint Csub using the connectivity map of a single network with a fairly high level of accuracy (≥80%), when Csub scans were pooled with short-term longitudinal scans from 26 HNU1 participants, and the Csub maps were drawn from the same site. In the HNU1 sample, the accuracy of the fingerprinting was over 95% on average in the frontoparietal and salience networks, close to what was reported using a connectome-based approach (Finn et al., 2015). Finn et al. (Finn and Constable, 2016) reported that certain networks comprised of nodes in the frontal, parietal, and temporal association cortices were the most discriminative for fingerprinting, which is consistent with our results (FPN and SAL had the highest fingerprinting accuracy in HNU1). These networks also exhibit the highest inter-individual variations in function (Mueller et al., 2013). When selecting pairs of scans from different sites or scanner vendors, fingerprinting accuracy of Csub fell to the lower end of the range of HNU1 distribution, with the exception of the visual network, where it was below the range. We thus observed a substantial impact of site and vendor effects on fingerprinting accuracy. Only one previous study had investigated multisite rsfMRI fingerprinting to our knowledge, and Hawco and colleagues (Hawco et al., 2018) had only included four subjects, and had not quantified the impact of site effect on fingerprinting accuracy, so their results cannot be directly compared to ours.

### 4.4 Study limitations

The major drawback of our study was its inclusion of only a single, male participant. This limitation is due to feasibility constraints, namely the time span of the study (scheduled to last at least five years) and the number of sites involved (set to increase to over 30 sites across Canada and other countries as CDIP is being rolled out to various recruiting sites in supported studies such as the CCNA). We report here on the first wave of data, collected over the initial 3.5 years. In order to assess that this single individual observation may be generalizable to other subjects, we confirmed that the intra-subject consistency we observed in Csub was close to what was observed on average in many HNU1 subjects (N=26) scanned 10 times over the course of one month at a single site. Our findings suggest that the consistency of connectivity maps remain of the same magnitude over several years, at least for a middle aged, healthy subject.

We also found that average FD differed between all three scanner vendors, and average tSNR and n_vols differed between Philips and Siemens scanners. The effect of n_vols ranged from 0.025 to −0.014, while the differences in consistency related to vendors were as large as 0.158 (Siemens, limbic network). The effect of n_vols observed accounted for differences in consistency for a difference of 72 seconds of scan time. While we cannot conclude firmly for longer acquisition times (20 minutes or more), by extrapolating the values we observe, the effect of n_vols when doubling or tripling the acquisition time would be as large as vendors effects. In other words, the consistency of 30 mns inter-vendor scans between vendors could get as high as the consistency of 10 mns intra-site scans. This conclusion is consistent with the results of Lauman et al. (Laumann et al., 2015), which sees a gain of over 0.1 in consistency when using 30 minutes of fMRI acquisition, rather than 10 minutes. Noble et al. (Noble et al., 2017a) also concluded that acquisition time may be as or even more important than site effects for reliability. These results are important to bear in mind when interpreting effects of scanner vendors on reproducibility of fMRI connectivity, as they appeared to be confounded by other important factors, at least in our sample, namely number of time samples, amount of motion and tSNR. Further steps should be taken in the future to further harmonize tSNR and motion in CDIP and other multisite protocols.

We also found that Csub was substantially less consistent with subjects from HNU1 than inter-subject consistency within HNU1. This observation may be due to the fact that HNU1 connectivity maps may be more similar because they were scanned at the same site. It also may reflect a difference in ethnicity between Csub (Caucasian) and HNU1 participants (a study based in China), and/or a systematic difference in age (Csub was older than HNU1 participants). These differences are a limitation of the data sample available for this study, and may have contributed to inflate the results of the fingerprinting experiment, with Csub reaching almost perfect fingerprinting accuracy in several networks when employing scans from the same site.

### 4.5 Alternate preprocessing

We did not evaluate the effects of field-map distortion correction on the consistency of rsfMRI measures. Recently, Togo and colleagues (Togo et al., 2017) reported improved detection of rsfMRI connectivity following field-map distortion correction on a 240 volume single-site dataset acquired on a 3T Siemens scanner. Connectivity was assessed with and without field-map distortion correction in several networks near the paranasal sinuses in the frontal lobe or the mastoid air cells and ear canals in the temporal lobe, brain regions most susceptible to distortion caused by magnetic field inhomogeneity (Jezzard and Balaban, 1995). A significant increase in connectivity strength was shown in the default-mode network, a network demonstrating robust consistency in our study. However, only a modest improvement in detection of the cerebellar network was reported, a network with lower consistency in our study (Togo et al., 2017). Moreover, we did not evaluate the effects of physiological noise correction on consistency, since the effect of this correction on consistently measures lacks consensus (Marchitelli et al., 2016).

## 5 CONCLUSIONS

### 5.1 Precision medicine

Despite negative effects of site and vendor differences, we still observed a fair level of consistency and fingerprinting accuracy of rsfMRI maps in our study, even for inter-vendor scans. It may thus be possible to extract multivariate biomarkers of brain diseases from such multisite harmonized data, as planned in the CCNA. Recent papers (Abraham et al., 2016; Orban et al., 2017) also showed that machine learning models trained on multisite rsfMRI data generalize better to subjects from new unseen sites, than models trained on single site data. Better approaches for site harmonization, either prospective like CDIP, or retrospective, for example (Yan et al., 2013), may still increase the precision of rsfMRI biomarkers, and is an important area of future work.

## 6. ACKNOWLEDGEMENTS

We would like to thank Dr. Bratislav Misic for suggesting the fingerprinting experiment, as well as the CCNA LORIS platform (Imaging, Database & Information Technology) for organizing the data, and Hanad Sharmarke for making available the Jupyter notebooks on binder. A.B. is currently supported by a Canadian Institute for Health Research (CIHR) Postdoctoral Fellowship (funding reference number #152548), CCNA, and the Courtois Foundation. At the start of the project A.B was supported by the Alzheimer Society of Canada Postdoctoral Fellowship. P.B. is supported by the CCNA and the Courtois Foundation. Financial support for I.C. and O.P. was obtained from the Alzheimer’s Society of Canada [grant number 13-32], the CIHR [grant number 117121], and the Fonds de recherche du Québec – Santé / Pfizer Canada - Pfizer-FRQS Innovation Fund [grant number 25262]. J.V. is supported by a Vanier Canada Graduate Studies Doctoral scholarship. S.D. is a Research Scholar from the Fonds de recherche du Québec – Santé [grant number 30801]. The *Consortium d’identification précoce de la maladie d’Alzheimer – Québec* is financed through the Fonds de recherche du Québec – Santé / Pfizer Canada Innovation Fund [grant number 25262]. The Canadian Consortium on Neurodegeneration in Aging is supported by a grant from the Canadian Institutes of Health Research with funding from several partners including the Alzheimer Society of Canada, Sanofi, and Women’s Brain Health Initiative.

## 7. SUPPLEMENTARY MATERIAL

Provided are Table S1, S2 and S3 referred to in Section 3.2 of the manuscript.

**Table S1:** Effect of each individual vendor and site for each of the seven rsfMRI networks. In this model vendor, site and time were used as explanatory variables. (*) indicated p<0.05, uncorrected for multiple comparisons.

|  | CER | DMN | FPN | LIM | MOT | SAL | VIS |
| --- | --- | --- | --- | --- | --- | --- | --- |
| CHUM | 0.1210 | *0.1822 | 0.1431 | *0.1964 | *0.1945 | 0.0955 | *0.1949 |
| CHUS | -0.0298 | -0.0257 | -0.0131 | -0.0665 | -0.0557 | -0.0246 | -0.0259 |
| CINQ | 0.0037 | -0.0087 | 0.0110 | -0.0366 | -0.0414 | -0.0171 | 0.0218 |
| ISMD | *0.1036 | -0.0193 | -0.0148 | 0.0617 | 0.0587 | *0.0912 | 0.0536 |
| IUGM | 0.0569 | -0.0027 | 0.0817 | 0.0218 | 0.0519 | 0.0677 | 0.0400 |
| MNI | *0.1536 | 0.0641 | *0.1085 | 0.0717 | 0.0887 | 0.0778 | 0.0169 |
| GE | 0.1412 | 0.0349 | 0.1470 | 0.0563 | 0.0011 | 0.0445 | 0.0099 |
| Philips | *0.0418 | -0.0048 | 0.0096 | 0.0291 | 0.0134 | *0.0458 | 0.0131 |
| Siemens | *0.0553 | *0.1091 | *0.0955 | *0.1507 | *0.0901 | *0.0811 | *0.1249 |

**Table S2:**
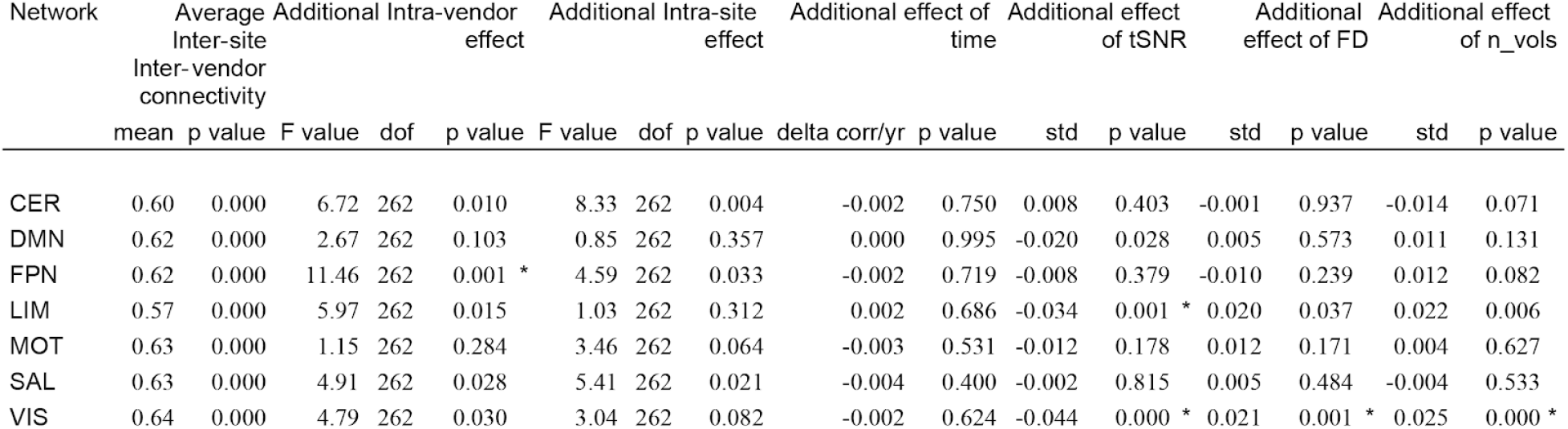
Effect of vendor, site, time, tSNR, FD, and n_vols on consistency of rsfMRI connectivity measures. The consistency of maps generated with rsfMRI data acquired on scanners from different vendors remained unchanged. As before, there was no substantial (or significant) effect of time between scanning sessions on consistency between maps. The estimated yearly rate of change in consistency ranged from 2.12e-3, p = 0.69 (limbic network) to −3.53e-3, p = 0.40 (salience network). There was a significant effect of vendors in the frontoparietal network, with trends (p<0.05 uncorrected) in four others (cerebellar, limbic, salience and visual). Intra-site consistency was not significantly higher than inter-site, inter-vendor consistency in any network, though trends (p<0.05 uncorrected) were observed in three networks (cerebellar, frontoparietal and salience). There was a significant effect of tSNR in the two networks (limbic and visual), with the default mode network indicating a trend (p<0.05 uncorrected). For both FD and n_vols, we found a significant effect in the visual network, with the limbic network indicating a trend (p<0.05 uncorrected). (*) indicates family-wise error < 0.05 (Bonferroni corrected for multiple comparisons across networks, adjusted threshold p<0.001). Abbreviations: Networks: CER, cerebellar; DMN, default mode; FPN, frontoparietal; LIM, limbic; MOT, motor; SAL, salience; VIS, visual.

**Table S3:** Effect of each individual vendor and site for each of the seven rsfMRI networks. In this model vendor, site, time, FD, n_vols and tSNR were used as explanatory variables. (*) indicated p<0.05, uncorrected for multiple comparisons.

|  | CER | DMN | FPN | LIM | MOT | SAL | VIS |
| --- | --- | --- | --- | --- | --- | --- | --- |
| CHUM | 0.1293 | *0.1648 | 0.1386 | 0.1690 | *0.1845 | 0.0944 | *0.1616 |
| CHUS | -0.0221 | -0.0428 | -0.0168 | -0.0991 | -0.0686 | -0.0271 | -0.0664 |
| CINQ | 0.0038 | -0.0088 | 0.0075 | -0.0330 | -0.0388 | -0.0157 | 0.0250 |
| ISMD | *0.1147 | -0.0321 | -0.0163 | 0.0291 | 0.0463 | *0.0898 | 0.0171 |
| IUGM | 0.0648 | -0.0044 | 0.0740 | 0.0173 | 0.0540 | 0.0716 | 0.0365 |
| MNI | *0.1408 | 0.0514 | *0.1050 | 0.0725 | 0.0858 | 0.0727 | 0.0048 |
| GE | 0.1425 | 0.0231 | 0.1474 | 0.0359 | -0.0088 | 0.0409 | -0.0172 |
| Philips | 0.0325 | -0.0062 | 0.0344 | 0.0308 | 0.0055 | *0.0378 | 0.0211 |
| Siemens | *0.0545 | *0.1087 | *0.0913 | *0.1557 | *0.0933 | *0.0824 | *0.1288 |

**Figure S1:**
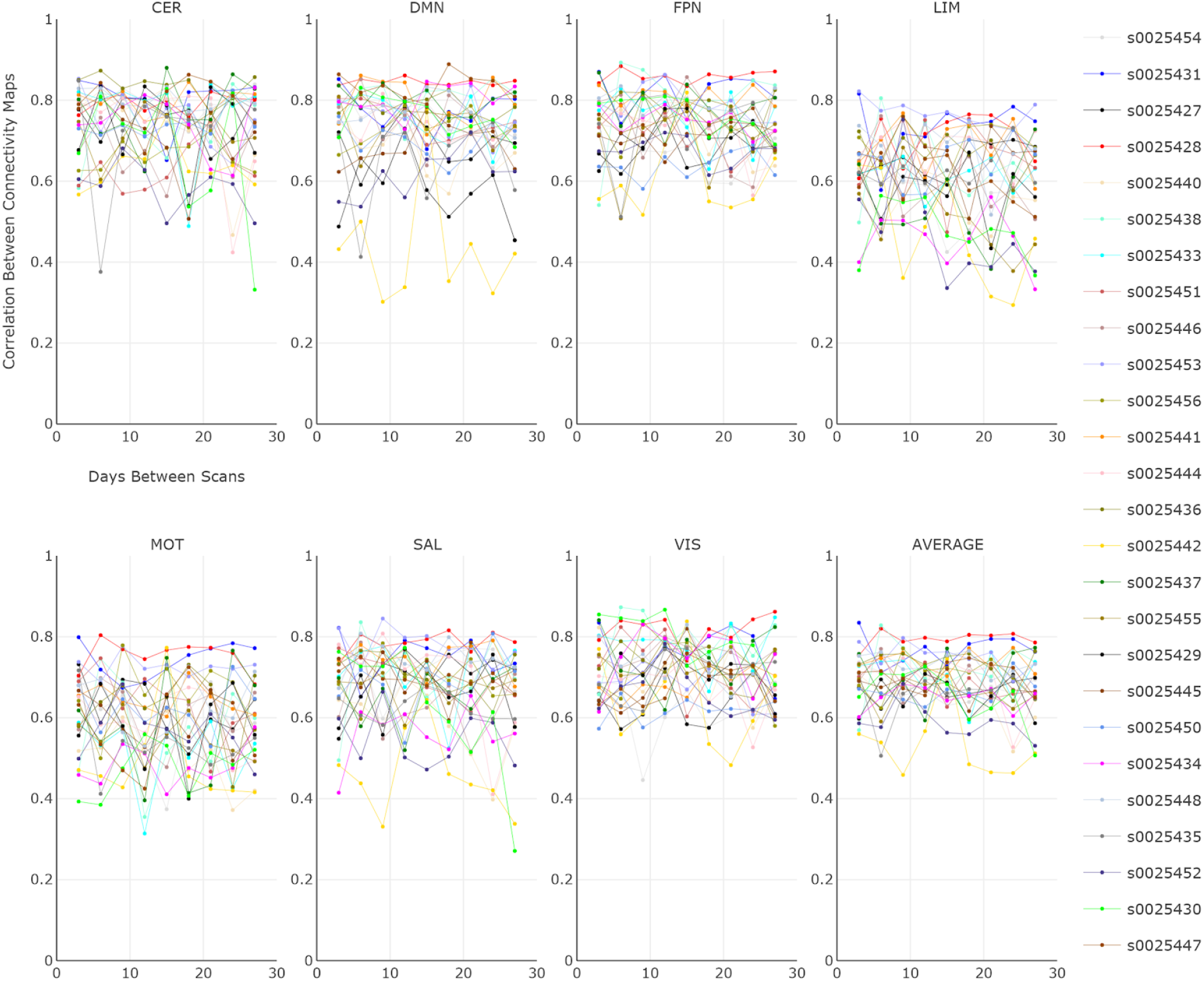
Intra- and inter-subject consistencies. Intra- and inter-subject consistency for the 26 individuals in the HNU1 dataset are plotted per network. The average consistency across all networks are also provided. To visualize the effect of time on intra-subject consistency, we plotted the consistency between connectivity maps at scan session 0 with that at scan sessions 1 to 9 (e.g. subject s0025438: scan session 0 vs scan session 1, subject s0025438: scan session 0 vs scan session 2). Per subject, the time interval between scan sessions was 3 days, with a total of 10 scans across one month. Abbreviations: CER, cerebellar network; DMN, default mode network; FPN, frontoparietal network; LIM, limbic network; MOT, motor network; SAL, salience network; VIS, visual network. Note: Interactive graphs are provided in the “graphs” Jupyter notebook.

## Footnotes

1 http://adni.loni.usc.edu/about/

2 http://www.ukbiobank.ac.uk/

3 https://addictionresearch.nih.gov/abcd-study

4 http://ccna-ccnv.ca/

5 www.cdip-pcid.ca

6 http://fcon_1000.projects.nitrc.org/indi/CoRR/html/hnu_1.html

7 http://fcon_1000.projects.nitrc.org/indi/CoRR/html/hnu_1.html

8 http://fcon_1000.projects.nitrc.org/indi/CoRR/html/_static/scan_parameters/HNU_1_scantable.pdf

9 https://hub.docker.com/r/simexp/niak-cog/

10 https://github.com/SIMEXP/niak/releases/download/v1.1.3/niak_singularity.tgz

11 http://www.gnu.org/software/octave/

12 http://www.bic.mni.mcgill.ca/ServicesSoftware/ServicesSoft-wareMincToolKit

13 http://mybinder.org

14 https://mybinder.org/v2/gh/SIMEXP/cdip_human_phantom/master

15 https://zenodo.org/record/1313201#.XAGdbVZK

16 https://plot.ly/

17 http://niak.simexp-lab.org/pipe_preprocessing.html

18 http://niak.simexp-lab.org/pipe_connectome.html

19 https://github.com/SIMEXP/cdip_human_phantom

20 https://doi.org/10.5281/zenodo.3350885

21 https://mybinder.org/v2/gh/SIMEXP/cdip_human_phantom/master

## REFERENCES

Abraham, A., Milham, M., Martino, A.D., Craddock, R.C., Samaras, D., Thirion, B., Varoquaux, G., 2016. Deriving reproducible biomarkers from multi-site resting-state data: An Autism-based example. Neuroimage. https://doi.org/10.1016/j.neuroimage.2016.10.045

Ad-Dab’bagh, Y., Lyttelton, O., Muehlboeck, J.S., Lepage, C., Einarson, D., Mok, K., Ivanov, O., Vincent, R.D., Lerch, J., Fombonne, E., Others, 2006. The CIVET image-processing environment: a fully automated comprehensive pipeline for anatomical neuroimaging research, in: Proceedings of the 12th Annual Meeting of the Organization for Human Brain Mapping. Florence, Italy, p. 2266.

An, H.S., Moon, W.-J., Ryu, J.-K., Park, J.Y., Yun, W.S., Choi, J.W., Jahng, G.-H., Park, J.-Y., 2017. Inter-vender and test-retest reliabilities of resting-state functional magnetic resonance imaging: Implications for multi-center imaging studies. Magn. Reson. Imaging 44, 125–130.

Badhwar, A., Tam, A., Dansereau, C., Orban, P., Hoffstaedter, F., Bellec, P., 2017. Resting-state network dysfunction in Alzheimer’s disease: A systematic review and meta-analysis. Alzheimers. Dement. 8, 73–85.

Bellec, P., Carbonell, F.M., Perlbarg, V., Lepage, C., Lyttelton, O., Fonov, V., Janke, A., Tohka, J., Evans, A.C., 2011. A neuroimaging analysis kit for Matlab and Octave, in: Proceedings of the 17th International Conference on Functional Mapping of the Human Brain. pp. 2735–2746.

Brown, M.R.G., Sidhu, G.S., Greiner, R., Asgarian, N., Bastani, M., Silverstone, P.H., Greenshaw, A.J., Dursun, S.M., 2012. ADHD-200 Global Competition: diagnosing ADHD using personal characteristic data can outperform resting state fMRI measurements. Front. Syst. Neurosci. 6, 69.

Button, K.S., Ioannidis, J.P.A., Mokrysz, C., Nosek, B.A., Flint, J., Robinson, E.S.J., Munafò, M.R., 2013. Power failure: why small sample size undermines the reliability of neuroscience. Nat. Rev. Neurosci. 14, 365.

Cheng, W., Palaniyappan, L., Li, M., Kendrick, K.M., Zhang, J., Luo, Q., Liu, Z., Yu, R., Deng, W., Wang, Q., Ma, X., Guo, W., Francis, S., Liddle, P., Mayer, A.R., Schumann, G., Li, T., Feng, J., 2015. Voxel-based, brain-wide association study of aberrant functional connectivity in schizophrenia implicates thalamocortical circuitry. NPJ Schizophr 1, 15016.

Cicchetti, D.V., Sparrow, S.A., 1981. Developing criteria for establishing interrater reliability of specific items: applications to assessment of adaptive behavior. Am. J. Ment. Defic. 86, 127–137.

Dansereau, C., Benhajali, Y., Risterucci, C., Pich, E.M., Orban, P., Arnold, D., Bellec, P., 2017. Statistical power and prediction accuracy in multisite resting-state fMRI connectivity. Neuroimage 149, 220–232.

Di Martino, A., O’Connor, D., Chen, B., Alaerts, K., Anderson, J.S., Assaf, M., Balsters, J.H., Baxter, L., Beggiato, A., Bernaerts, S., Blanken, L.M.E., Bookheimer, S.Y., Braden, B.B., Byrge, L., Castellanos, F.X., Dapretto, M., Delorme, R., Fair, D.A., Fishman, I., Fitzgerald, J., Gallagher, L., Keehn, R.J.J., Kennedy, D.P., Lainhart, J.E., Luna, B., Mostofsky, S.H., Müller, R.-A., Nebel, M.B., Nigg, J.T., O’Hearn, K., Solomon, M., Toro, R., Vaidya, C.J., Wenderoth, N., White, T., Craddock, R.C., Lord, C., Leventhal, B., Milham, M.P., 2017. Enhancing studies of the connectome in autism using the autism brain imaging data exchange II. Sci Data 4, 170010.

Dong, A., Toledo, J.B., Honnorat, N., Doshi, J., Varol, E., Sotiras, A., Wolk, D., Trojanowski, J.Q., Davatzikos, C., Alzheimer’s Disease Neuroimaging Initiative, 2017. Heterogeneity of neuroanatomical patterns in prodromal Alzheimer’s disease: links to cognition, progression and biomarkers. Brain 140, 735–747.

Drysdale, A.T., Grosenick, L., Downar, J., Dunlop, K., Mansouri, F., Meng, Y., Fetcho, R.N., Zebley, B., Oathes, D.J., Etkin, A., Schatzberg, A.F., Sudheimer, K., Keller, J., Mayberg, H.S., Gunning, F.M., Alexopoulos, G.S., Fox, M.D., Pascual-Leone, A., Voss, H.U., Casey, B.J., Dubin, M.J., Liston, C., 2017. Resting-state connectivity biomarkers define neurophysiological subtypes of depression. Nat. Med. 23, 28–38.

Finn, E.S., Shen, X., Scheinost, D., Rosenberg, M.D., Huang, J., Chun, M.M., Papademetris, X., Constable, R.T., 2015. Functional connectome fingerprinting: identifying individuals using patterns of brain connectivity. Nat. Neurosci. 18, 1664–1671.

Finn, E.S., Todd Constable, R., 2016. Individual variation in functional brain connectivity: implications for personalized approaches to psychiatric disease. Dialogues Clin. Neurosci. 18, 277–287.

Fleiss, J.L., Cohen, J., 1973. The Equivalence of Weighted Kappa and the Intraclass Correlation Coefficient as Measures of Reliability. Educ. Psychol. Meas. 33, 613–619.

Friedman, L., Glover, G.H., 2006. Report on a multicenter fMRI quality assurance protocol. J. Magn. Reson. Imaging 23, 827–839.

Friedman, L., Glover, G.H., Krenz, D., Magnotta, V., FIRST BIRN, 2006. Reducing inter-scanner variability of activation in a multicenter fMRI study: role of smoothness equalization. Neuroimage 32, 1656–1668.

Giove, F., Gili, T., Iacovella, V., Macaluso, E., Maraviglia, B., 2009. Images-based suppression of unwanted global signals in resting-state functional connectivity studies. Magn. Reson. Imaging 27, 1058–1064.

Gordon, E.M., Laumann, T.O., Gilmore, A.W., Newbold, D.J., Greene, D.J., Berg, J.J., Ortega, M., Hoyt-Drazen, C., Gratton, C., Sun, H., Hampton, J.M., Coalson, R.S., Nguyen, A.L., McDermott, K.B., Shimony, J.S., Snyder, A.Z., Schlaggar, B.L., Petersen, S.E., Nelson, S.M., Dosenbach, N.U.F., 2017. Precision Functional Mapping of Individual Human Brains. Neuron 95, 791–807.e7.

Gratton, C., Laumann, T.O., Nielsen, A.N., Greene, D.J., Gordon, E.M., Gilmore, A.W., Nelson, S.M., Coalson, R.S., Snyder, A.Z., Schlaggar, B.L., Dosenbach, N.U.F., Petersen, S.E., 2018. Functional Brain Networks Are Dominated by Stable Group and Individual Factors, Not Cognitive or Daily Variation. Neuron 98, 439–452.e5.

Hawco, C., Viviano, J.D., Chavez, S., Dickie, E.W., Calarco, N., Kochunov, P., Argyelan, M., Turner, J.A., Malhotra, A.K., Buchanan, R.W., Voineskos, A.N., SPINS Group, 2018. A longitudinal human phantom reliability study of multi-center T1-weighted, DTI, and resting state fMRI data. Psychiatry Res Neuroimaging. https://doi.org/10.1016/j.pscychresns.2018.06.004

Hunter, J.D., 2007. Matplotlib: A 2D Graphics Environment. Comput. Sci. Eng. 9, 90–95.

Jezzard, P., Balaban, R.S., 1995. Correction for geometric distortion in echo planar images from B0 field variations. Magn. Reson. Med. 34, 65–73.

Jones, E., Oliphant, T., Peterson, P., 2016. others. SciPy: Open source scientific tools for Python. 2001. URL http://www.scipy.org.

Jovicich, J., Minati, L., Marizzoni, M., Marchitelli, R., Sala-Llonch, R., Bartrés-Faz, D., Arnold, J., Benninghoff, J., Fiedler, U., Roccatagliata, L., Picco, A., Nobili, F., Blin, O., Bombois, S., Lopes, R., Bordet, R., Sein, J., Ranjeva, J.-P., Didic, M., Gros-Dagnac, H., Payoux, P., Zoccatelli, G., Alessandrini, F., Beltramello, A., Bargalló, N., Ferretti, A., Caulo, M., Aiello, M., Cavaliere, C., Soricelli, A., Parnetti, L., Tarducci, R., Floridi, P., Tsolaki, M., Constantinidis, M., Drevelegas, A., Rossini, P.M., Marra, C., Schönknecht, P., Hensch, T., Hoffmann, K.-T., Kuijer, J.P., Visser, P.J., Barkhof, F., Frisoni, G.B., PharmaCog Consortium, 2016. Longitudinal reproducibility of default-mode network connectivity in healthy elderly participants: A multicentric resting-state fMRI study. Neuroimage 124, 442–454.

Koo, T.K., Li, M.Y., 2016. A Guideline of Selecting and Reporting Intraclass Correlation Coefficients for Reliability Research. J. Chiropr. Med. 15, 155–163.

Laumann, T.O., Gordon, E.M., Adeyemo, B., Snyder, A.Z., Joo, S.J., Chen, M.-Y., Gilmore, A.W., McDermott, K.B., Nelson, S.M., Dosenbach, N.U.F., Schlaggar, B.L., Mumford, J.A., Poldrack, R.A., Petersen, S.E., 2015. Functional System and Areal Organization of a Highly Sampled Individual Human Brain. Neuron 87, 657–670.

Li, H., Satterthwaite, T.D., Fan, Y., 2018. BRAIN AGE PREDICTION BASED ON RESTING-STATE FUNCTIONAL CONNECTIVITY PATTERNS USING CONVOLUTIONAL NEURAL NETWORKS. Proc. IEEE Int. Symp. Biomed. Imaging 2018, 101–104.

Marchitelli, R., Minati, L., Marizzoni, M., Bosch, B., Bartrés-Faz, D., Müller, B.W., Wiltfang, J., Fiedler, U., Roccatagliata, L., Picco, A., Nobili, F., Blin, O., Bombois, S., Lopes, R., Bordet, R., Sein, J., Ranjeva, J.-P., Didic, M., Gros-Dagnac, H., Payoux, P., Zoccatelli, G., Alessandrini, F., Beltramello, A., Bargalló, N., Ferretti, A., Caulo, M., Aiello, M., Cavaliere, C., Soricelli, A., Parnetti, L., Tarducci, R., Floridi, P., Tsolaki, M., Constantinidis, M., Drevelegas, A., Rossini, P.M., Marra, C., Schönknecht, P., Hensch, T., Hoffmann, K.-T., Kuijer, J.P., Visser, P.J., Barkhof, F., Frisoni, G.B., Jovicich, J., 2016. Test-retest reliability of the default mode network in a multi-centric fMRI study of healthy elderly: Effects of data-driven physiological noise correction techniques. Hum. Brain Mapp. 37, 2114–2132.

Matthews, P.M., Hampshire, A., 2016. Clinical Concepts Emerging from fMRI Functional Connectomics. Neuron 91, 511–528.

McKinney, W., Others, 2010. Data structures for statistical computing in python, in: Proceedings of the 9th Python in Science Conference. Austin, TX, pp. 51–56.

Mueller, S., Wang, D., Fox, M.D., Yeo, B.T.T., Sepulcre, J., Sabuncu, M.R., Shafee, R., Lu, J., Liu, H., 2013. Individual variability in functional connectivity architecture of the human brain. Neuron 77, 586–595.

Nielsen, J.A., Zielinski, B.A., Fletcher, P.T., Alexander, A.L., Lange, N., Bigler, E.D., Lainhart, J.E., Anderson, J.S., 2013. Multisite functional connectivity MRI classification of autism: ABIDE results. Front. Hum. Neurosci. 7, 599.

Noble, S., Scheinost, D., Finn, E.S., Shen, X., Papademetris, X., McEwen, S.C., Bearden, C.E., Addington, J., Goodyear, B., Cadenhead, K.S., Mirzakhanian, H., Cornblatt, B.A., Olvet, D.M., Mathalon, D.H., McGlashan, T.H., Perkins, D.O., Belger, A., Seidman, L.J., Thermenos, H., Tsuang, M.T., van Erp, T.G.M., Walker, E.F., Hamann, S., Woods, S.W., Cannon, T.D., Constable, R.T., 2017a. Multisite reliability of MR-based functional connectivity. Neuroimage 146, 959–970.

Noble, S., Spann, M.N., Tokoglu, F., Shen, X., Constable, R.T., Scheinost, D., 2017b. Influences on the Test-Retest Reliability of Functional Connectivity MRI and its Relationship with Behavioral Utility. Cereb. Cortex 27, 5415–5429.

Oliphant, T.E., 2006. A guide to NumPy. Trelgol Publishing USA.

Orban, P., Dansereau, C., Desbois, L., Mongeau-Pérusse, V., Giguère, C.-É., Nguyen, H., Mendrek, A., Stip, E., Bellec, P., 2017. Multisite generalizability of schizophrenia diagnosis classification based on functional brain connectivity. Schizophr. Res. https://doi.org/10.1016/j.schres.2017.05.027

Pedregosa, F., Varoquaux, G., Gramfort, A., Michel, V., Thirion, B., Grisel, O., Blondel, M., Prettenhofer, P., Weiss, R., Dubourg, V., Vanderplas, J., Passos, A., Cournapeau, D., Brucher, M., Perrot, M., Duchesnay, É., 2011. Scikit-learn: Machine Learning in Python. J. Mach. Learn. Res. 12, 2825–2830.

Power, J.D., Barnes, K.A., Snyder, A.Z., Schlaggar, B.L., Petersen, S.E., 2012. Spurious but systematic correlations in functional connectivity MRI networks arise from subject motion. Neuroimage 59, 2142–2154.

Saggar, M., Tsalikian, E., Mauras, N., Mazaika, P., White, N.H., Weinzimer, S., Buckingham, B., Hershey, T., Reiss, A.L., Diabetes Research in Children Network (DirecNet), 2017. Compensatory Hyperconnectivity in Developing Brains of Young Children With Type 1 Diabetes. Diabetes 66, 754–762.

Seabold, S., Perktold, J., 2010. Statsmodels: Econometric and statistical modeling with python, in: Proceedings of the 9th Python in Science Conference. SciPy society Austin, p. 61.

Seeley, W.W., Crawford, R.K., Zhou, J., Miller, B.L., Greicius, M.D., 2009. Neurodegenerative diseases target large-scale human brain networks. Neuron 62, 42–52.

Skåtun, K.C., Kaufmann, T., Doan, N.T., Alnæs, D., Córdova-Palomera, A., Jönsson, E.G., Fatouros-Bergman, H., Flyckt, L., KaSP, Melle, I., Andreassen, O.A., Agartz, I., Westlye, L.T., 2017. Consistent Functional Connectivity Alterations in Schizophrenia Spectrum Disorder: A Multisite Study. Schizophr. Bull. 43, 914–924.

Togo, H., Rokicki, J., Yoshinaga, K., Hisatsune, T., Matsuda, H., Haga, N., Hanakawa, T., 2017. Effects of Field-Map Distortion Correction on Resting State Functional Connectivity MRI. Front. Neurosci. 11, 656.

Urchs, S., Armoza, J., Benhajali, Y., St-Aubin, J., Orban, P., Bellec, P., 2017. MIST: A multi-resolution parcellation of functional brain networks. MNI Open Res 1, 3.

Van Dijk, K.R.A., Sabuncu, M.R., Buckner, R.L., 2012. The influence of head motion on intrinsic functional connectivity MRI. Neuroimage 59, 431–438.

Yan, C.-G., Craddock, R.C., Zuo, X.-N., Zang, Y.-F., Milham, M.P., 2013. Standardizing the intrinsic brain: towards robust measurement of inter-individual variation in 1000 functional connectomes. Neuroimage 80, 246–262.

Yeo, B.T.T., Krienen, F.M., Sepulcre, J., Sabuncu, M.R., Lashkari, D., Hollinshead, M., Roffman, J.L., Smoller, J.W., Zöllei, L., Polimeni, J.R., Fischl, B., Liu, H., Buckner, R.L., 2011. The organization of the human cerebral cortex estimated by intrinsic functional connectivity. J. Neurophysiol. 106, 1125–1165.

Zuo, X.-N., Anderson, J.S., Bellec, P., Birn, R.M., Biswal, B.B., Blautzik, J., Breitner, J.C.S., Buckner, R.L., Calhoun, V.D., Castellanos, F.X., Chen, A., Chen, B., Chen, J., Chen, X., Colcombe, S.J., Courtney, W., Craddock, R.C., Di Martino, A., Dong, H.-M., Fu, X., Gong, Q., Gorgolewski, K.J., Han, Y., He, Y., He, Y., Ho, E., Holmes, A., Hou, X.-H., Huckins, J., Jiang, T., Jiang, Y., Kelley, W., Kelly, C., King, M., LaConte, S.M., Lainhart, J.E., Lei, X., Li, H.-J., Li, K., Li, K., Lin, Q., Liu, D., Liu, J., Liu, X., Liu, Y., Lu, G., Lu, J., Luna, B., Luo, J., Lurie, D., Mao, Y., Margulies, D.S., Mayer, A.R., Meindl, T., Meyerand, M.E., Nan, W., Nielsen, J.A., O’Connor, D., Paulsen, D., Prabhakaran, V., Qi, Z., Qiu, J., Shao, C., Shehzad, Z., Tang, W., Villringer, A., Wang, H., Wang, K., Wei, D., Wei, G.-X., Weng, X.-C., Wu, X., Xu, T., Yang, N., Yang, Z., Zang, Y.-F., Zhang, L., Zhang, Q., Zhang, Z., Zhang, Z., Zhao, K., Zhen, Z., Zhou, Y., Zhu, X.-T., Milham, M.P., 2014. An open science resource for establishing reliability and reproducibility in functional connectomics. Sci Data 1, 140049.

